# Microgravity effect on murine T cells exposed to suborbital flight aboard Blue Origin’s New Shepard vehicle

**DOI:** 10.1101/2021.05.13.443970

**Authors:** Pedro J. Llanos, Kristina Andrijauskaite, Vijay Vishal Duraisamy, Sathya Gangadharan, Jay Morris, Michael J. Wargovich

## Abstract

Numerous scientific experiments have been conducted in space. However, the precise mechanisms mediating successful human body adaption to the hostile space environment are still not delineated. The cost and logistic challenges of sending biological payloads to International Space Station are forcing scientists to find alternative research platforms. In this study, we investigated whether brief exposure to microgravity during the suborbital flight aboard Blue Origin’s New Shepard rocket modulated the behavior of the gravity-sensitive murine T cells. We assessed the effect of suborbital environment on different T cell subsets, activation markers, functionality, and cytokine secretion capabilities. Thus, to optimize the potential response of T cells, we cultured them in interleukin IL-2 alone or combined with IL-12. We found that exposure to microgravity decreased the expression of T cells with CD4+ cells being more sensitive to suborbital flight as compared to CD8+ cells. Our data indicate that the functional capabilities of flown T cells were reduced. Also, our findings suggest that supplementing cells with IL-2 and IL-12 cytokines may restore microgravity-mediated cellular alterations. Finally, our study provides insights on the microgravity effect on the murine T cells by utilizing a novel suborbital research platform.

## Introduction

There has been an escalated interest over the past few years by NASA and other private stakeholders to pursue human spaceflight missions to the moon (e.g., ARTEMIS mission) and beyond [1], [2], [3]. However, space is a hostile environment associated with numerous health hazards, including dysregulation of the immune system [4], [5], [6], [2]. The body of research conducted in this field is quite substantial, but with conflicting findings and unclear mechanisms on how spaceflight induces immune system alterations. Therefore, more research is needed to facilitate more definite conclusions and to set up basis for interventions [7]. Research findings derived from sampling astronauts and conducting in-vitro and in-vivo experiments reveal that spaceflight affects several immune parameters including reduced production of cytokines, decreased proliferation rates, and suppressed functional capabilities of different types of immune cells [8]. However, several studies conducted in the 90s showed that spaceflight enhanced the production of certain cytokines [9], [10] and had conflicting effects on T cell subpopulations with several studies demonstrating a decreased expression of CD4+ (helper cytokine production) and CD8+ (cytotoxic activity) cells [11], [12] while others showing an increased expression of CD4+ cells post flight [13]. This may be attributed to methodological alterations, such as the use of different stimulants (e.g., ConA, phytohemagglutinin, PMA/I), diverse animal models, and the specific flight profiles of different space missions. One of the mechanisms leading to dysregulated immune system during or after space travel is susceptibility to infections and viruses [14]. The main defense machinery to fight infections is through T lymphocytes (T cells). They coordinate the host response against microbial and cancerous developments leading to elimination and long-term protection [15]. At first, naive T lymphocytes need to be activated so that they can differentiate into effector cells to perform their immune functions. This requires engagement of TCR (T Cell Receptor) by the peptide/major histocompatibility complex (MHC) and the costimulatory signal provided by the interaction between accessory molecules, such as CD28 and CD3 on the surface of T cells [16]. Upon activation, T cells proliferate and differentiate into effector T cells driven by autocrine interleukin-2 (IL-2) and other cytokines. T cells treated with various concentrations of interleukins IL-2 and IL-12 can have different responses for mediating inflammation and other infectious events. The ideal environment to study microgravity induced alterations on the immune system is the International Space Station (ISS). However, because of the cost and long launch preparation times, novel research platforms exist to enhance our understanding of the impact of microgravity on immunity. Those platforms include terrestrial spaceflight simulations, parabolic flights, balloons, clinostats, rotating wall vessels, random position machines, etc. However, it is necessary to distinguish the extent to which these models mimic microgravity. Currently, there are a very few accessible U.S. national space research platforms (Blue Origin’s New Shepard, Virgin Galactic’s Space Ship Two, Exos’s SARGE-1 rocket from Aerospace Systems and Technologies) providing yearly launches with a quick turnaround payload recovery and high quality continuous microgravity time of about 3-5 minutes depending of the flight provider and respective rocket flight profile (∼100-120 km) as well as other European vehicles (MASER 14 – suborbital express) that can reach apogee altitude of about 240 km providing approximately 6 minutes of microgravity. These different companies and their vehicles give scientists opportunities to fly payloads at various apogees and diverse microgravity exposure times [17]. See table 1.

**Table 1:** Different microgravity research platforms.

| Platform | Microgravity level (g) | Duration | Volume (m3) | Waiting Time | Cost/Experiment (\$) |
| --- | --- | --- | --- | --- | --- |
| Free fall towers | $10^{-3} - 10^{-6}$ | < 5 s | < 1 | months | 5 k\$ |
| Parabolic flights | $10^{-2} - 10^{-3}$ | ~ 20-25 s | > 10 | Months -1 year | 125 k\$ |
| Sounding rockets | $10^{-4} - 10^{-5}$ | 5-13 min | < 1 | > 2 years | > 400 k\$ |
| ISS | $10^{-2} - 10^{-5}$ | weeks, months to years | > 1 | > 5 years | 1-5 M\$ |
| Blue Origin's New Shepard | $10^{-2} - 10^{-3}$ | 3.2-3.4 min | NanoLab (0.5 kg) (10cmx10cmx20cm) | 6 months to 1year | ~8,000-16,000 |
| Blue Origin's New Shepard | $10^{-2} - 10^{-3}$ | 3.2-3.4 min | 0.0489 (11.34 kg) | 6 months to 1year | ≥100 k\$ |
| Exos' SARGE -1 | ~0.005 | 3-4 min | 1U (10cmx10cmx10cm) to 2U | 6 months to 1 year | ~ 5-10 k\$ |
| MASER-14 | $\leq 10^{-4}$ | ~ 6 min | <1 (10cmx10cmx20cm) | 1 year to 2 years | ≥ 25 k\$ |

Thus, suborbital flights provide researchers the opportunity to design experiments, refine hypotheses and collect preliminary data, which could be further tested using orbital platforms after having matured the technologies. Furthermore, this unique research platform allows for a more controlled experimental design because of its larger flexibility and shorter sample retrieval time. In this study, we investigated the influence of altered gravity on murine T cells during the suborbital mission aboard Blue Origin’s New Shepard rocket. Given T cells are sensitive to environmental changes and acknowledging microgravity induced immune changes demonstrated by other researchers, we hypothesized that even a brief exposure to microgravity during the suborbital space flight would lead to alterations of T cell behavior and function. Further, we added IL-2 and IL-12 alone or combined at different time-points to the T cell cultures. Our findings demonstrate that even a brief exposure of about 3.2 minutes to microgravity modulated the activity of T cells as measured by the alterations in different T cell subsets, cell surface markers’ expression, functional, and cytokine secretion capabilities.

## Materials and Methods

### Splenocytes Isolation and T cell Activation

Spleens were dissected from euthanized C57BL/6 healthy mice, derived from the breeding colony (2013044AR) which were maintained in UT Health Science Center (UTHSCSA) at specific pathogen-free facility according to the Institutional Animal Use and Care Committee (IACUC) standards. T cells were separated from the already euthanized animals derived from the breeding colony with the following UT Health IACUC committee approved protocol #2013044AR. Euthanasia was administered in a bell jar with isoflurane as the inhalation for induction or as a route of isoflurane administration.

Spleens from female mice (n = 3) with age of 6-8 months were used to set up primary cultures. Single cell-suspensions of lymphocytes were achieved by sterile dissociation of whole spleens with syringe plunger through cell strainer and by lysing red blood cells with the ACK (Ammonium-Chloride-Potassium) lysing buffer (Life Technologies). Isolated splenocytes were pooled together and placed in 24-well plates at 10^6^ cells per ml concentration in 1.5 ml and activated with anti-CD3 mAb (145-2C11 clone, plate-bound, 1 µg/ml) and anti-CD28 mAb (37.51 clone, soluble, 2 µg/ml) from Bio X Cell in RPMI media for 48 hours with or without human (h) IL-2 (200 ng/ml, recombinant human interleukin-2 (rIL-2)) from NCI at Frederick Repository. After 48 h, cells were washed twice and supplemented either with no cytokines or with human (h) IL-2 (200 ng/ml) and mouse (m) mIL-12 (10 ng/ml) from Shenandoah Biotechnology alone or in combination every three days.

### Transportation of T cells and maintenance at West Texas Launch Facility

T cells were transported during a six-hour drive from UT Health to Van Horn (Texas) to the Payload Processing Facility (PPF) in West Texas Launch Site (WTLS). Detailed description of the transportation procedures and conditions at PPF and WTLS were described in a previous manuscript [18]. Briefly, T cells were kept at 34 °C to 35 °C during transportation and in a self-made incubator assembled (L-3 days) from the heated box with the tubing attached to the life support system containing 5% carbon dioxide, 21% oxygen, and the rest was balanced with nitrogen. At L-2, 1 days, T cells were kept at about 25.5 °C during the day and at about 28.0 °C overnight in the Fisher Scientific electrical heater. The launch was scrubbed for one day.

### Flight Conditions and Ground Controls

Initial estimation of the payload weight allowed the housing of nine 5 mL Eppendorf tubes in the NanoLab including (in duplicate): T cells with no cytokines, T cells with IL-2, T cells with IL-12, T cells with IL-2/IL-12, and T cells primed with IL-2 since day 0, during activation process (not in duplicate). The same conditions pertained to the ground controls. After preliminary payload integration into the NanoLab, the team estimated to have additional space and therefore included three additional tubes containing cells supplemented with an extra dose of IL-2, IL-12, and IL-2/IL-12 added one hour before handing off the payload to the NanoRacks team or about 10 hours before the flight. Those conditions lacked ground controls and were referred as IL-2, IL-12, IL-2/IL-12 before flight.

The diagram below (diagram 1) illustrates the different timeline when cytokines were added.

**Diagram 1:** Diagram of T cell culture supplementation with different cytokines.

| Condition | During 48 h<br>initial Activation | Timeline post 48 h |
| --- | --- | --- |
| **Flight:** |  |  |
| no cytokines (0) | - | - |
| IL-2 | - | continuously |
| IL-2 day 0 onward | IL-2 | continuously |
| IL-12 | - | continuously |
| IL-2/-IL-12 | - | continuously |
| IL-2 before flight | - | continuously & before flight |
| IL-12 before flight | - | continuously & before flight |
| IL-2/IL-12 before flight | - | continuously & before flight |
| **Ground:** |  |  |
| no cytokines (0) | - | - |
| IL-2 | - | continuously |
| IL-2 day 0 onward | IL-2 | continuously |
| IL-12 | - | continuously |
| IL-2/IL-12 | - | continuously |

### Flow Cytometry

Cells were analyzed by flow cytometry using standard procedures as previously described [19]. Briefly, cells were washed in staining buffer (PBS, 2% bovine growth serum and 0.01% sodium azide) and stained with fluorescently labelled antibodies. The fluorescently conjugated monoclonal antibodies (mAb) used in this study included: anti-mouse (m) CD4 mAb (RM4-5, dilution 1/500), anti-mCD279 (PD-1) mAb (J43, dilution 1/250) from eBiosience; anti-mCD8 mAb (53-6.7, dilution 1/500), anti-mCD25 mAb (PC61, dilution 1/250), anti-mIFNγ mAb (XMG1.2, dilution 1/250), anti-mCD71 mAb (RI7217, dilution 1/250) and anti-mTNFα mAb (MP6-XT722, dilution 1/250) from Biolegend; anti-mCD69 mAb (H1.2F3, dilution 1/250) from BD Biosciences. For intracellular cytokine expression, we used the BD Cytofix/Cytoperm Kit (BD Biosciences, San Jose, CA, USA). Samples were acquired on Celesta and data were analyzed using the FloJo software (Tree Star, Ashland, OR, US).

### FITC Annexin V/Dead Cell Apoptosis Assay

T cells were stained with FITC Anenxin V/PI (Invitrogen) following manufacturer’s instructions and different apoptosis stages were determined by flow cytometry. The amount of total apoptosis (%) was calculated by taking the percentage of cells positive for early and late apoptotic markers.

### T cell Functional Assays

To assay the ability of T cells to respond functionally to antigen, we cultured cells for 6 h with either PMA (100 ng/ml, Sigma), Ionomycin (1 µM, Sigma), and Golgistop (BD) or with hTyr peptide (1 µg/ml) (YMDGTMSQV, American Peptide Company). Cultures were set up with in a flat-bottom 96-well plates. After incubation at 37 °C, cells were stained for intracellular cytokines and assayed by flow cytometry.

### Multiplex Immunoassay

Cell culture supernatants were collected from flown T cells and ground controls and stored at - 80 °C until further processing. The supernatants were analyzed simultaneously for 5 cytokines, including mouse IL-12, IL-2, IL-4, IL-6, and IL-10 employing a bead-based sandwich immunoassay. For the detection of multiple cytokines, a monoclonal antibody specific for each cytokine was coupled to a particular set of beads with known internal fluorescence. Cytokine secretion assay was performed using ThermoFisher ProcartaPlex Multiplex Custom Made mouse cytokine 5-plex assay kit following manufacturer’s instructions and analyzed with the Bio-Plex 200 Luminex-based multiplex analysis system (Bio-Rad, Hercules, CA). All samples were analyzed using technical duplicates. The expression of different cytokines was determined by the fluorescence intensity (FI).

### Statistical Analysis

Data were graphically displayed using prism 6 software. Differences in various markers expressions across different conditions were determined and compared using unpaired t-tests with a two-sided alpha of 0.05/0.01 or a two-way ANOVA. Data are represented as mean ± standard error mean (SEM) or ± standard deviation (SD).

## Results

### Description of the Blue Origin suborbital M7 mission, sample recovery and processing

Launching this scientific payload entitled Cell Research Experiment in Microgravity (CRExIM) aboard Blue Origin’s New Shepard Mission 7 was part of the multidisciplinary project between different United States universities as a collaborative effort to advance the knowledge of microgravity induced T cell alterations. Blue Origin’s New Shepard is the reusable suborbital rocket designed to fly prospective astronauts and research payloads past the Kármán line. It is a vertical takeoff and landing (VTVL) space vehicle. The system consists of an updated pressurized capsule, New Shepard 2.0, atop a booster. It accelerates for approximately two and a half minutes before the main engine cut off. The capsule then separates from the booster to coast into suborbital space. After one minute of free fall, the booster performs an autonomously controlled rocket-powered vertical landing, while the capsule lands under parachutes after about 10 minutes of flight. For the M7 mission with our payload aboard, the New Shepard vehicle reached a maximum altitude of about 322,440 feet (98.3 km), and experienced microgravity for about 3.2 minutes, landing within one mile of the launch pad after takeoff (Fig 1).

**Fig 1.**
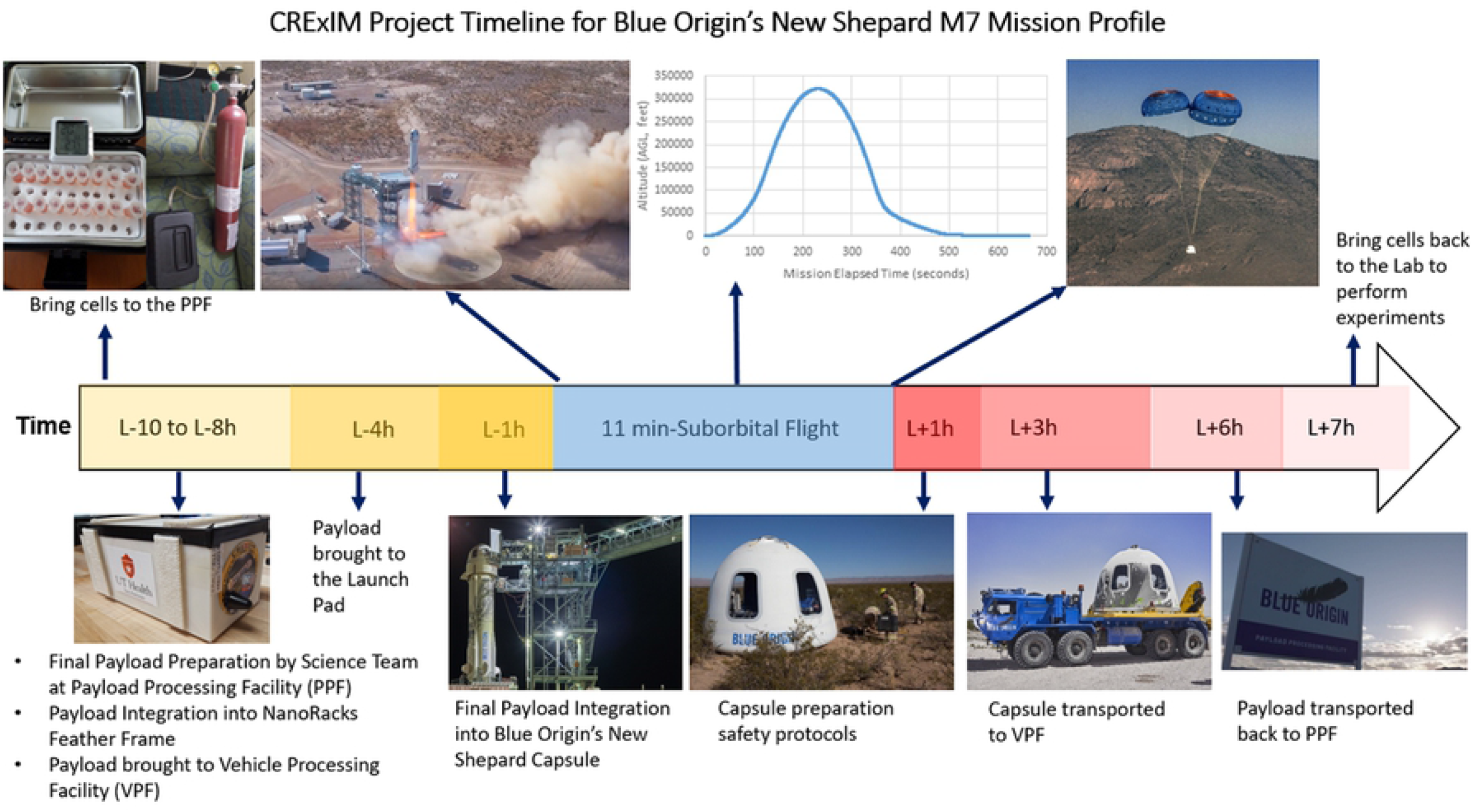
Representation of the M7 mission profile and corresponding pre-flight and post-flight procedures. From left to right indicate the timeline (hours) and the activity/procedures taken in regard to the suborbital launch (L). Minus (-) indicates prior to the launch, while (+) indicates post-launch time point.

The NanoLab was recovered 6 hours after liftoff. The temperature inside the NanoLab was tracked during pre-flight (6-8 hours before liftoff), flight (10 minutes 10 seconds), and post-flight operations (6 hours after liftoff and before being delivered to our team). While in the Payload Processing Facility (PPF), T cells were kept at 27 °C before integration into the vehicle. The reason to keep T cells at this temperature and not at 37 °C, which is the standard temperature for culturing cells, was to avoid exposing the T cells to a temperature shock given the freezing conditions in the desert at West Texas with temperatures as low as 2 °C to -4 °C at night. Also, we knew that the thermal conditions inside the capsule were going to be near 20 °C as communicated by the Blue Origin team in their payload’s guide [20]. A thermal sensor was placed inside one of the sides of the NanoLab while the 5 mL Eppendorf tubes that contained the T cells were inside a foam structure to protect them from structural vibrations and as a layer of containment. The temperature of the flown cells was about 23 °C during the 10-minute suborbital flight and recovered 6 hours later at about 25 °C [18].

### Microgravity alters expression of CD8+ and especially CD4+ T cell subsets

To investigate whether brief exposure to microgravity leads to alterations in CD8+ and CD4+ subsets, we stained cells with fluorescently labelled antibodies for these markers. Table 2 shows the cell population for CD4+, CD8+, total apoptosis, live cells, and dead cells and their corresponding means, mean differences, standard error (SE), and the relative error (RE) of the means for both control (ground) and flight conditions indicating that the RE is higher for CD4+ than for CD+8 subsets. Consistent with findings from other groups [21], our data also indicate that exposure to microgravity leads to altered expression of CD8 and CD4 positive cells (Figs 2A, 2B).

**Table 2:** T cell Population Analysis of Means.

| Cell Population | Ground Mean | Flight Mean | Diff ± SE | Relative Error Means |
| --- | --- | --- | --- | --- |
| CD4+ | 75.57 | 70.27 | 5.3 ± 0.5 | 7.0 |
| CD8+ | 64.3 | 62.9 | 1.4 ± 0.3 | 2.1 |
| Total Apoptosis | 35.0 | 39.2 | -4.2 ± 1.0 | 12 |
| Live Cells | 48.1 | 50.2 | -2.1 ± 0.1 | 4.4 |
| Dead Cells | 16.1 | 10.6 | 5.5 ± 0.1 | 34.2 |

**Fig 2.**
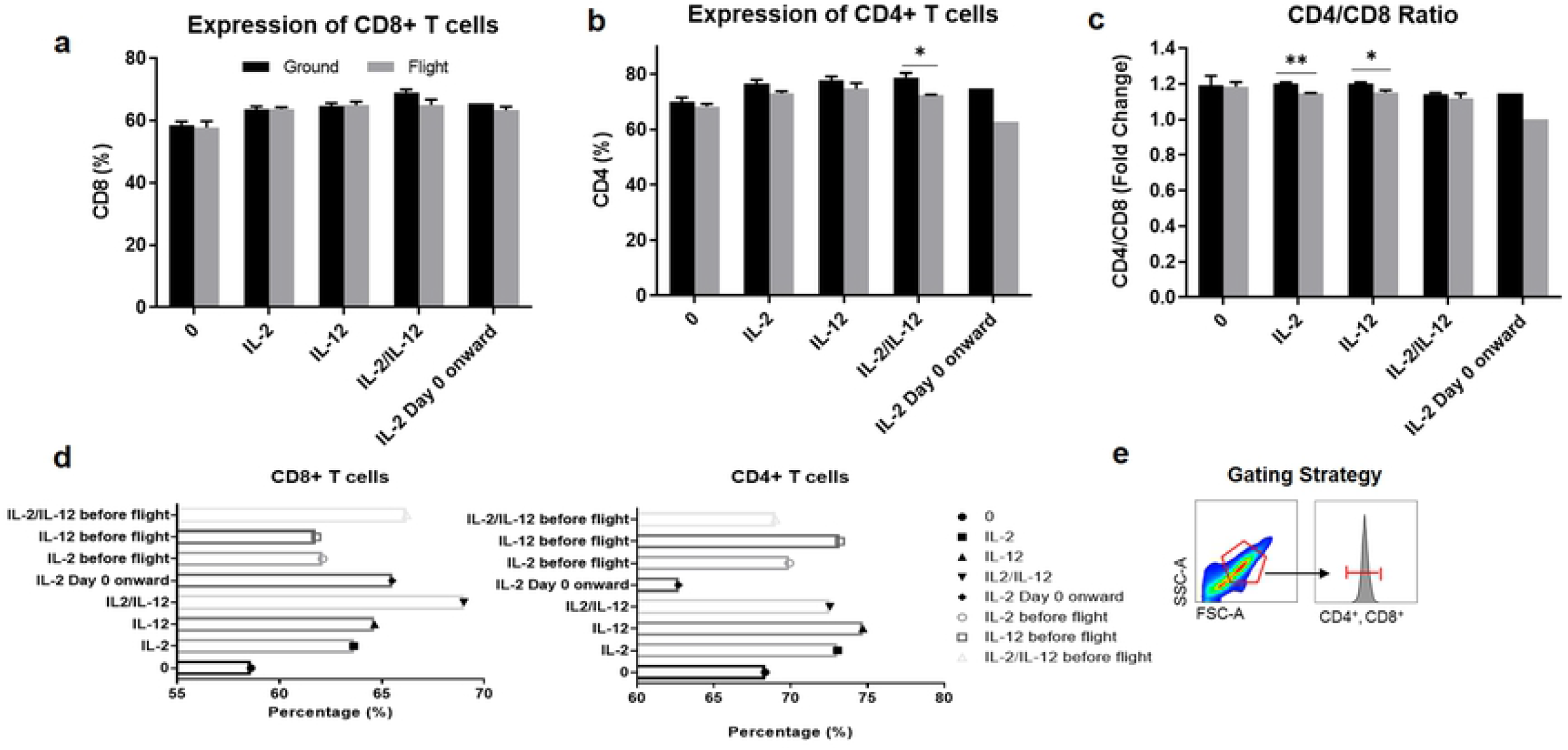
Brief exposure to microgravity decreases the expression of CD4+ and CD8+ subsets. Cells were assessed for CD8+ and CD4+ expression (%) by staining cells with fluorescently labeled antibodies for CD8 and CD4 by flow cytometry. (a) Indicates CD8+ and (b) CD4+ expression in culturing cells with or without cytokines (condition 0) across ground and flight conditions. (c) Displays CD4/CD8 subpopulation ratio in both flight and ground conditions. (d) Shows expression in additional flight conditions. (d) Gating strategy included drawing a gate around live cell population based on the FSC-A/SSC-A plot followed by the histograms of fluorescence peak heights for CD4+ and CD8+ expression. Data represented as means and means ±SEM. ** p<0.01, * p<0.05. Significance was determined by using t-tests. Black color represents ground conditions, while grey color indicates flight conditions.

More specifically, we found a slight decrease in CD8+ and CD4+ populations in all flight conditions with the most significant difference observed in cells cultured in both cytokines (p<0.05) (Fig 2B). Our data revealed that T cells exposed to suborbital flight had a decreased CD4+/CD8+ ratio in all conditions with the most profound differences observed in cells cultured in IL-2 (p<0.01) and IL-12 (p<0.05) (Fig 2C). This suggests that culturing T cells with cytokines have influence on the distribution of CD4+ and CD8+ expression. There was almost no variation in the ratios of CD4+ to CD8+ when no cytokines were added. The latter is consistent with some previous studies [22] which reported no difference between the CD4+ and CD8+ ratios. However, there was a profound change (near 12.2. % decrease) of this ratio when the IL-2 cytokine was supplied from day zero between the ground and flight conditions. Given we did not have this condition in duplicate and only one tube of cells was exposed to flight, the difference of expression did not reach significance. Finally, adding cytokines 10 hours prior to suborbital flight led to decreased expression of CD8+ CD4+ cell populations. Thus, our data indicate that CD4+ subset of cells cultured in IL-2 from day 0 throughout the whole duration of the experiment decreased the expression of CD4 positive cells when compared to all other conditions (Fig 2D). In alignment with previous reports [23], our results indicate that CD4+ subset might be more sensitive to microgravity exposure than CD8+ cells.

Column 1 shows the cell population for CD4+, CD8+, total apoptosis, live cells, and dead cells. Column 2 corresponds to the means of the ground samples for each of the previous populations. Column 3 depicts the means of the flight samples for each of the previous populations. Column 4 displays the difference between the means of control and flight conditions and its associated standard error (SE). Column 5 depicts the relative error (RE) of the means for both control (theoretical) and flight (measured) conditions indicating that the RE is higher for CD4+ than for CD+8 subsets. Numbers represent means (%) of indicated T cell populations. Data were obtained using fluorescence-labeled monoclonal antibodies and flow cytometry. Ground indicates ground conditions including cells cultured in the following: no cytokines, IL-2, IL-12, IL-2/IL-12, IL-2 from day 0 onward; flight indicates flight conditions cultured in the same manner as ground conditions. One-way ANOVA test was used to compute the SE and RE means.

### Microgravity decreases the expression of the T cell activation marker CD71 and T cell exhaustion marker PD-1

We next sought to determine whether T cells exposed to suborbital flight condition would have a decreased expression of early (CD69) mid-late (CD71) and late (CD25) activation markers as well as the alteration in the checkpoint programmed death-1 PD-1 marker, also known as T cell exhaustion indicator (Fig 3).

**Fig 3.**
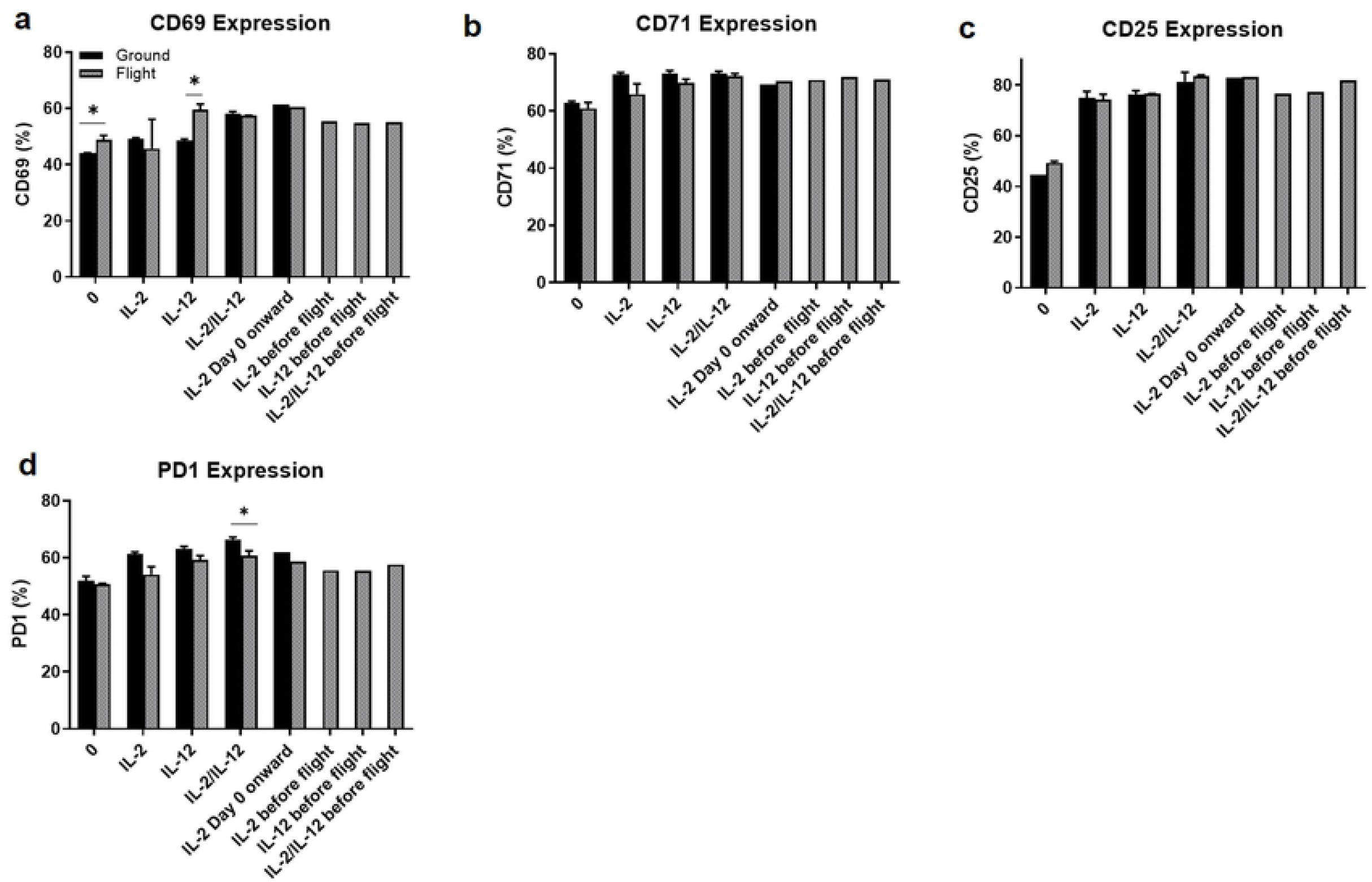
T cells exposed to microgravity exhibit either decreased or maintain similar expression of activation markers, except for cells cultured in IL-12, which have elevated levels of CD69. Flight conditions show a decreased expression of exhaustion marker PD1. T cells were stimulated with PMA and ionomycin for 6 hours. Then cells were washed and stained with fluorescently labeled indicated markers and assessed by flow cytometry. The percentage for CD69 (a), CD71 (b), CD25 (c), and PD1 (d) was determined by gating on live cells. (%). Data represented as means ±SEM. * p<0.05. Significance was determined by using the T-test. Black color represents ground conditions, while grey shows flight conditions.

T cell activation plays a significant role in cellular immune responses. Activated T cells express a variety of surface markers and they secrete Th1 cytokines interleukin-2 (IL-2) and interferon-γ (IFNγ). Although most previous microgravity related studies demonstrated the suppressed expression of activation markers in both rodents and humans, many studies could not rule out the influence of other confounders, such as cosmic radiation, variability in mission profiles and experimental methodologies [23].

Our data indicate that for CD69, flight cells cultured in IL-2 had a 7.2 % decrease, whereas flight cells cultured in IL-12 had a significant (p<0.05) increase in CD69 expression (Fig 3A). Thus, adding IL-2 before the flight also increased the expression of CD69 by about 21% when compared to IL-2 flight condition. CD69 is a classical early marker of lymphocyte activation due to its rapid appearance on the surface of the plasma membrane after stimulation. Given that CD69 regulates the secretion of IFNɣ whose expression is also influenced by the IL-12 cytokine, it is not surprising to see elevation of this marker in IL-12 condition. This suggests that culturing T cells in IL-12 might be a good strategy to increase the expression of CD69. However, it is interesting to observe this effect in flight condition only, but not in ground control. Previous studies indicated that the duration of the mission played a significant role in CD69 expression which was elevated in short duration shuttle crewmembers, but it was significantly reduced in long duration mission crewmembers [24].

CD71 expression across flight conditions were slightly reduced except for cells cultured in IL-2 from day 0 onward (Fig 3B). Thus, adding extra cytokines before the flight increased the expression of CD71 in IL-2 (7.5 %) and IL-12 (2.8 %) flight conditions (Fig 3B). This suggests that supplementing cells with these cytokines prior to the flight may restore the expression of CD71. CD25 expression remained similar among different flight and ground conditions with a slight increase (3%) in cells supplemented with additional IL-2 dose before the flight (Fig 3C). Since CD71 expression was downregulated in most flight conditions, it may be that this marker is more sensitive to microgravity than CD69 and CD25 markers. In addition, we observed that cells not cultured in cytokines led to the increase of about 11% of CD69 and CD25 activation markers, and a slight decrease of less than 3% for CD71 marker. Our data agree with findings from other researchers which showed the decrease expression of activation markers in flight conditions [15] and reduced proliferation capacity of lymphocytes [25]. Our experiments complement previous findings with additional insights on how culturing T cells using different cytokines affects the expression of activation markers, especially after supplementing cells with extra doses of cytokines just prior the suborbital flight. Differences in T cell activation markers observed across different conditions could indicate incomplete or inefficient function of signals required for full T cell activation or a longer exposure time needed to detect more significant changes.

Finally, we assessed the effect of microgravity on PD-1 expression (Fig 3D). PD-1 mediated-inhibitory signals play a major role in T cell exhaustion during chronic infections and cancers, which makes PD-1 a valuable target of checkpoint blockade in cancer immunotherapy. Given that T cells become exhausted in large tumors, we hypothesized that T cells could also become exhausted under microgravity conditions and this marker could be used as an indication of decreased T cell activity in microgravity studies. However, PD-1 is also considered an activation marker of CD4+ and CD8+ T cells and similar to CTLA-4 (cytotoxic T-lymphocyte-associated protein 4), may be upregulated early to potentially prime negative regulatory feedback mechanisms to limit inflammation [26]. Our results indicate that PD-1 expression was decreased in all conditions exposed to suborbital flight as compared to ground controls with the most significant effect in cells cultured in IL2/IL-12 (p<0.05) (Fig 3D). Furthermore, in agreement with the literature [27] which suggests that many cytokines can up-regulate PD-1 expression, we also observed this increased expression in all ground and flight conditions as compared to cells with no cytokines. There is some evidence to draw parallels between T cells in microgravity and in cancer. As such, our data demonstrated that T cells exposed to microgravity resulted in decreased expression of PD-1 whose expression is proportional to the strength of TCR signaling to compensate T cell activation and to control immune response [28]. Thus, it is also well-established that PD-1 expression on antigen-specific T cells reflects the functional avidity and anti-tumor reactivity of these cells [28]. Therefore, the importance of PD-1 on human immunity makes this molecule a valuable marker to explore its role in microgravity for immunotherapy purposes, especially given that slow immune response in space could put astronauts at risk for developing various infectious or accelerate the process of cancerous lesions. Given our observation of decreased expression of this marker across all flight conditions, this indicates that PD-1 might be sensitive to exposure to microgravity and could be used for future studies to examine T cell exhaustion under microgravity conditions. However, future studies should assess the expression of PD-1 at the protein level as well as after longer exposure to microgravity as suborbital flight might be too short period to detect the exhaustion of T cells.

### Microgravity drives cells to more apoptotic phenotype but decreases the expression of dead cells in flight conditions. Thus, culturing cells in IL-2 decreases the levels of apoptosis

After observing that brief exposure to microgravity can affect T cell markers of activation, we next investigated the levels of apoptosis (Table 2, Fig 4).

**Fig 4.**
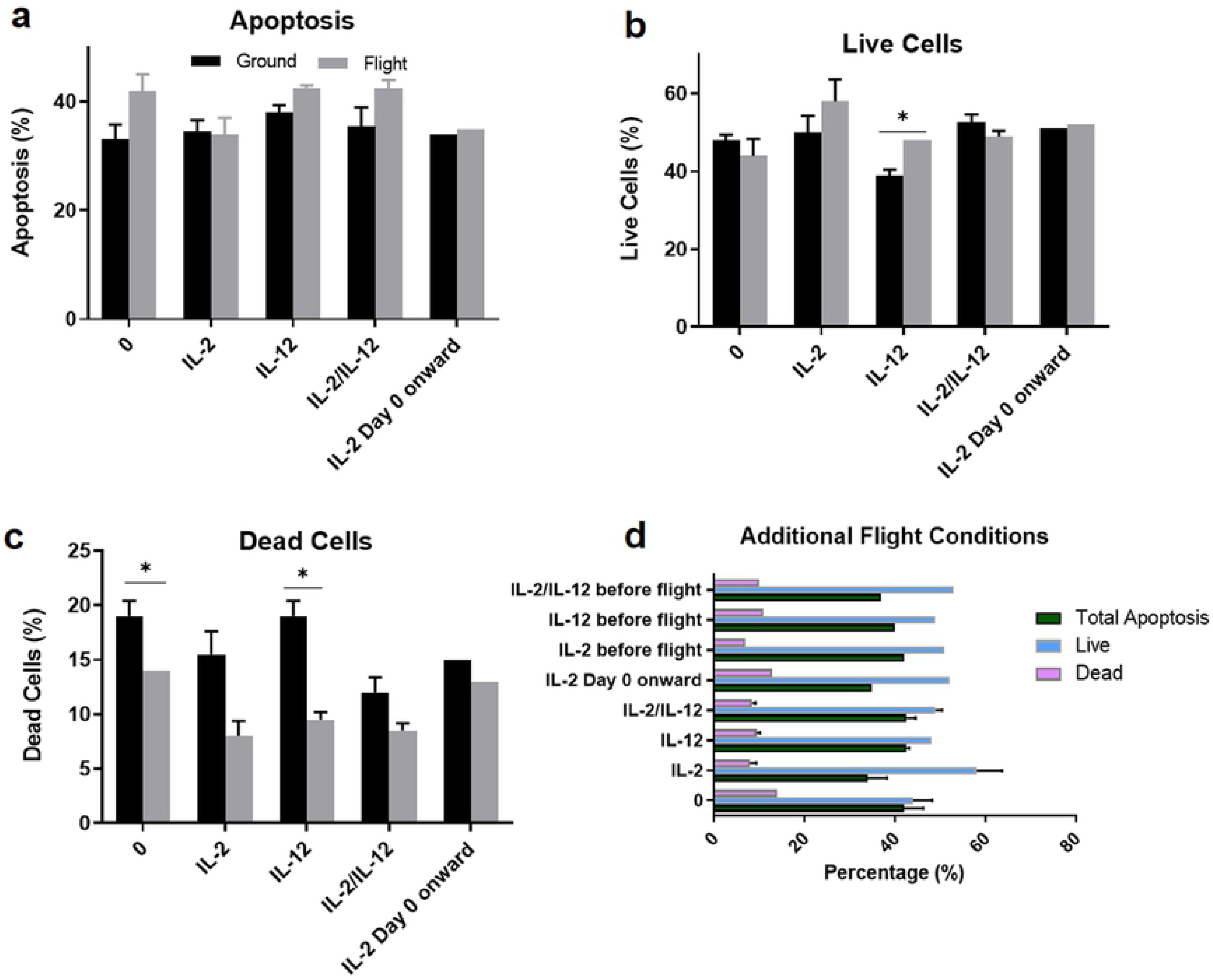
Brief exposure to microgravity drives cells to more apoptotic phenotype. However, culturing cells in different cytokines profoundly decreases the percentage of dead cells in flight conditions. Rates of total apoptosis (%) including early and late (a), live (b), dead (c), and additional flight conditions (d) were determined by staining cells with Annexin-V/PI and performing flow cytometric analysis. Columns indicate T cell culture conditions, while rows correspond to ground (top), flight (middle) and additional flight (bottom) conditions. Data represented as means ±SD. * p<0.05. Black color represents ground conditions, while grey color indicates flight conditions; green color shows the percentage of total apoptosis, light blue – live cells, while pink – dead cells.

We stained cells with AnnexinV/PI dyes and assessed the percentage of total apoptosis (early and late), as well as live and dead cell populations. Propidium iodide (PI) is a nucleic acid binding dye which is impermeable to live cells and apoptotic cells, but stains dead cells, binding tightly to the nucleic acids in the cells. Combining PI with Annexin V FITC antibody allows a clear distinction of different phases of apoptosis where apoptotic cells should express high levels of Annexin V, whereas dead cells should have high expression of both PI and Annexin V with live cells showing little or no fluorescence. Our data revealed that brief exposure to microgravity led to the increased levels of cell death in all conditions except for the cells primed with IL-2, with the most profound effect observed in cells with no cytokines (27%) and cells cultured in both cytokines (20%) (Fig 4A). Although IL-12 is a promising candidate for cancer immunotherapy, it is however, associated with some level of toxicity [29] which is reflected in our study by an increased level of apoptosis. Given that IL-12 may induce toxicity, our data also revealed that the highest apoptosis rate was observed in conditions with IL-12 alone or in combination with IL-2 (Fig 4A). Furthermore, our data showed that these conditions also had higher expression of PD-1 (Fig 3D) suggesting the correlation between PD-1 and apoptosis [30]. It has been reported in the literature [31] that up-regulation of PD-1 could promote T cell apoptosis in certain types of cancer. In addition, our results indicate that adding IL-2 and IL-12 to flown T cell cultures increases the percentage of live cells (Fig 4B). Compared to ground controls, flown cells had fewer dead cells in all flight conditions (Fig 4C). This suggests that microgravity may drive T cells into more apoptotic phenotype but may save them from dying. Notably, adding cytokines before the flight did not decrease the apoptosis nor increased the number of live cells, except in cells primed with IL-2 which resulted in decrease number of dead cells (Fig 4D). This suggests that IL-2 cytokine may be a good candidate to increase the live cells populations under microgravity exposure. Microgravity induced apoptosis was reported previously [32] in space flown lymphocytes (Jurkat).

### Microgravity leads to decreased functional ability of T cells as measured by decreased expression of IFNɣ and TNFα

To determine if brief exposure to microgravity affects the functional ability of T cells, we stimulated them with either PMA/I (Fig 5) or hTyr peptide (data not shown) for 6 hours and assessed their functional capabilities by staining for intracellular IFNγ and TNFα expression.

**Fig 5.**
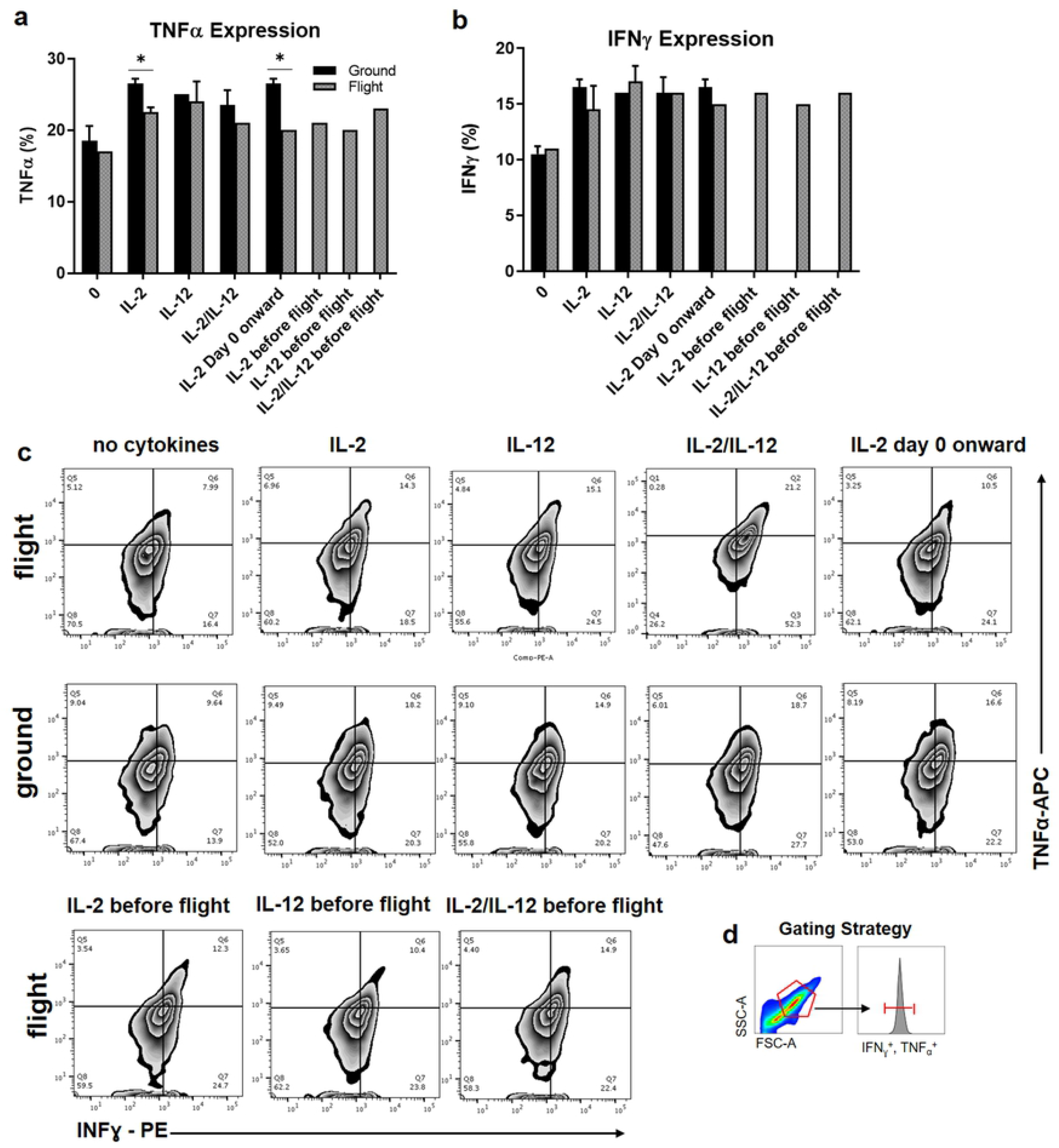
Brief exposure to microgravity leads to decreased expression of TNFα and IFNɣ, except for T cells cultured in IL-12, which exhibit greater levels of IFNɣ. Cells were incubated with PMA (100 ng/ml)/Ionomycin (1 µM) for 6h. Afterwards they were washed and stained for intracellular TNFα (a) and IFNɣ (b) expression. Green color represents ground conditions, blue color indicates flight conditions. Data represented as means ±SD. (c) Shows graphical representation of functional assessment of T cells after stimulation with hTyr peptide for 6 hours with expression of IFNɣ-PE (x axis) and TNFα-APC (y axis). Columns indicate T cell culture conditions, while rows correspond to flight (top), ground (middle) and additional flight (bottom) conditions. (d) Gating strategy included drawing a gate around live cell population based on the FSC-A/SSC-A plot followed by the histograms of fluorescence peak heights for IFNɣ + and TNFα+ expression.

Our data revealed that stimulation with cytokines led to increased expression of these two markers in all conditions (Figs 5A, 5B) as compared to no stimulation. However, there were differences observed across different cytokines used. There was a reduction in TNFα levels in all flight conditions when compared to ground controls, especially in cells treated with IL-2 (p<0.05) either regularly or continuously from day 0 (Fig 5A). However, there was an increase (10%) in TNFα additional IL-2/IL-12 cytokines were added before the flight in comparison to cells cultured in these cytokines continuously. For the IFNγ expression, there was about 6.3 % increase in flown cells cultured in IL-12 and a 10% increase in cells supplemented with extra dose of IL-2 just before the flight (Fig 5B). Culturing cells with IL-12 alone led to overall the highest expression of TNFα and IFNɣ among all flight conditions. This suggests that using this cytokine may elicit T helper (h) cells Th1 response, which is characterized by the generation of large amounts of IFNγ and TNFα. However, when compared flight IL-12 condition to ground control, there was a decrease in TNFα expression (about 4%), while we found an increase in IFNɣ expression (about 6%). Given that one of the roles of IL-12 cytokine is to induce IFNɣ, it is not surprising that cells cultured in IL-12 had the highest expression of IFNγ in flight conditions. A central mediator of IL-12-induced responses is IFNγ, which is secreted upon IL-12 stimulation alone or with synergizing factors such as IL-2 and IL-18. We stimulated cells with both (PMA/I) (Figs 5C, 5D) and peptide (data not shown). We did not observe a profound difference between these two stimulants suggesting that using either of these methods yields consistent results.

### Exposure to microgravity alters the expression of multiple cytokines detected in T cell culture supernatants

We collected supernatants from cells flown aboard Blue Origin’s New Shepard as well as from ground controls to perform cytokine secretion analysis using Multiplex Immunoassay kit. This extremely sensitive assay was chosen to detect the expression of five analytes in our T cell cultures including IL-2, IL-4, IL-6, IL-10, and IL-12 (Table 3).

**Table 3:**
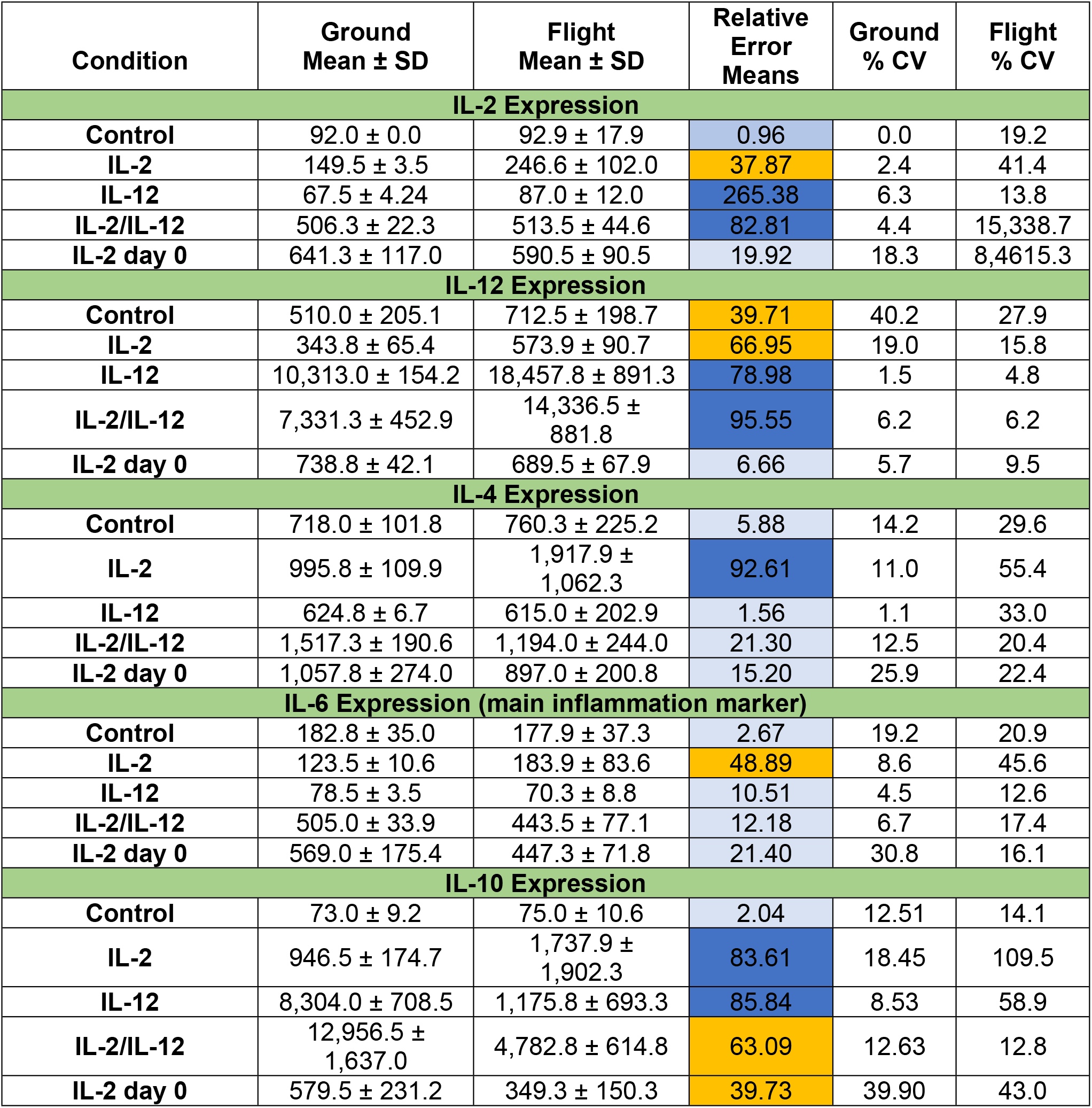
Cytokine Expression Analysis from T Cell Culture Supernatants.

| Condition | Ground Mean ± SD | Flight Mean ± SD | Relative Error Means | Ground % CV | Flight % CV |
| --- | --- | --- | --- | --- | --- |
| **IL-2 Expression** |  |  |  |  |  |
| Control | 92.0 ± 0.0 | 92.9 ± 17.9 | 0.96 | 0.0 | 19.2 |
| IL-2 | 149.5 ± 3.5 | 246.6 ± 102.0 | 37.87 | 2.4 | 41.4 |
| IL-12 | 67.5 ± 4.24 | 87.0 ± 12.0 | 265.38 | 6.3 | 13.8 |
| IL-2/IL-12 | 506.3 ± 22.3 | 513.5 ± 44.6 | 82.81 | 4.4 | 15,338.7 |
| IL-2 day 0 | 641.3 ± 117.0 | 590.5 ± 90.5 | 19.92 | 18.3 | 8,4615.3 |
| **IL-12 Expression** |  |  |  |  |  |
| Control | 510.0 ± 205.1 | 712.5 ± 198.7 | 39.71 | 40.2 | 27.9 |
| IL-2 | 343.8 ± 65.4 | 573.9 ± 90.7 | 66.95 | 19.0 | 15.8 |
| IL-12 | 10,313.0 ± 154.2 | 18,457.8 ± 891.3 | 78.98 | 1.5 | 4.8 |
| IL-2/IL-12 | 7,331.3 ± 452.9 | 14,336.5 ± 881.8 | 95.55 | 6.2 | 6.2 |
| IL-2 day 0 | 738.8 ± 42.1 | 689.5 ± 67.9 | 6.66 | 5.7 | 9.5 |
| **IL-4 Expression** |  |  |  |  |  |
| Control | 718.0 ± 101.8 | 760.3 ± 225.2 | 5.88 | 14.2 | 29.6 |
| IL-2 | 995.8 ± 109.9 | 1,917.9 ± 1,062.3 | 92.61 | 11.0 | 55.4 |
| IL-12 | 624.8 ± 6.7 | 615.0 ± 202.9 | 1.56 | 1.1 | 33.0 |
| IL-2/IL-12 | 1,517.3 ± 190.6 | 1,194.0 ± 244.0 | 21.30 | 12.5 | 20.4 |
| IL-2 day 0 | 1,057.8 ± 274.0 | 897.0 ± 200.8 | 15.20 | 25.9 | 22.4 |
| **IL-6 Expression (main inflammation marker)** |  |  |  |  |  |
| Control | 182.8 ± 35.0 | 177.9 ± 37.3 | 2.67 | 19.2 | 20.9 |
| IL-2 | 123.5 ± 10.6 | 183.9 ± 83.6 | 48.89 | 8.6 | 45.6 |
| IL-12 | 78.5 ± 3.5 | 70.3 ± 8.8 | 10.51 | 4.5 | 12.6 |
| IL-2/IL-12 | 505.0 ± 33.9 | 443.5 ± 77.1 | 12.18 | 6.7 | 17.4 |
| IL-2 day 0 | 569.0 ± 175.4 | 447.3 ± 71.8 | 21.40 | 30.8 | 16.1 |
| **IL-10 Expression** |  |  |  |  |  |
| Control | 73.0 ± 9.2 | 75.0 ± 10.6 | 2.04 | 12.51 | 14.1 |
| IL-2 | 946.5 ± 174.7 | 1,737.9 ± 1,902.3 | 83.61 | 18.45 | 109.5 |
| IL-12 | 8,304.0 ± 708.5 | 1,175.8 ± 693.3 | 85.84 | 8.53 | 58.9 |
| IL-2/IL-12 | 12,956.5 ± 1,637.0 | 4,782.8 ± 614.8 | 63.09 | 12.63 | 12.8 |
| IL-2 day 0 | 579.5 ± 231.2 | 349.3 ± 150.3 | 39.73 | 39.90 | 43.0 |

Table 3 has 6 columns, the first column is the condition (control, IL-2, IL-12, IL-2 and IL-12 combined, IL-2 from day 0), the second and third columns display the means of the ground and flight samples with their corresponding standard deviations (SD), respectively; the fourth column depicts the relative error (RE) of ground and flight samples; and the fifth and sixth columns are the percentage of the coefficient of variability (CV) for ground and flight samples, respectively. Table 2 shows the absolute value of the relative error (RE) across different conditions. Small differences with a RE of 0-33% are shown in light blue, RE between 33% and 66% are shown in orange, and large RE differences greater than 66% are shown in dark blue. Data were obtained from T cell culture supernatants from flown T cells (flight) and their matched ground controls (ground) after performing ProcartaPlex Multiplex Custom-Made mouse cytokine 5-plex immunoassay analysis.

Our findings indicate that adding cytokines to T cell cultures lead to altered expression of many cytokines among different ground and flight conditions (Fig 6A).

**Fig 6:**
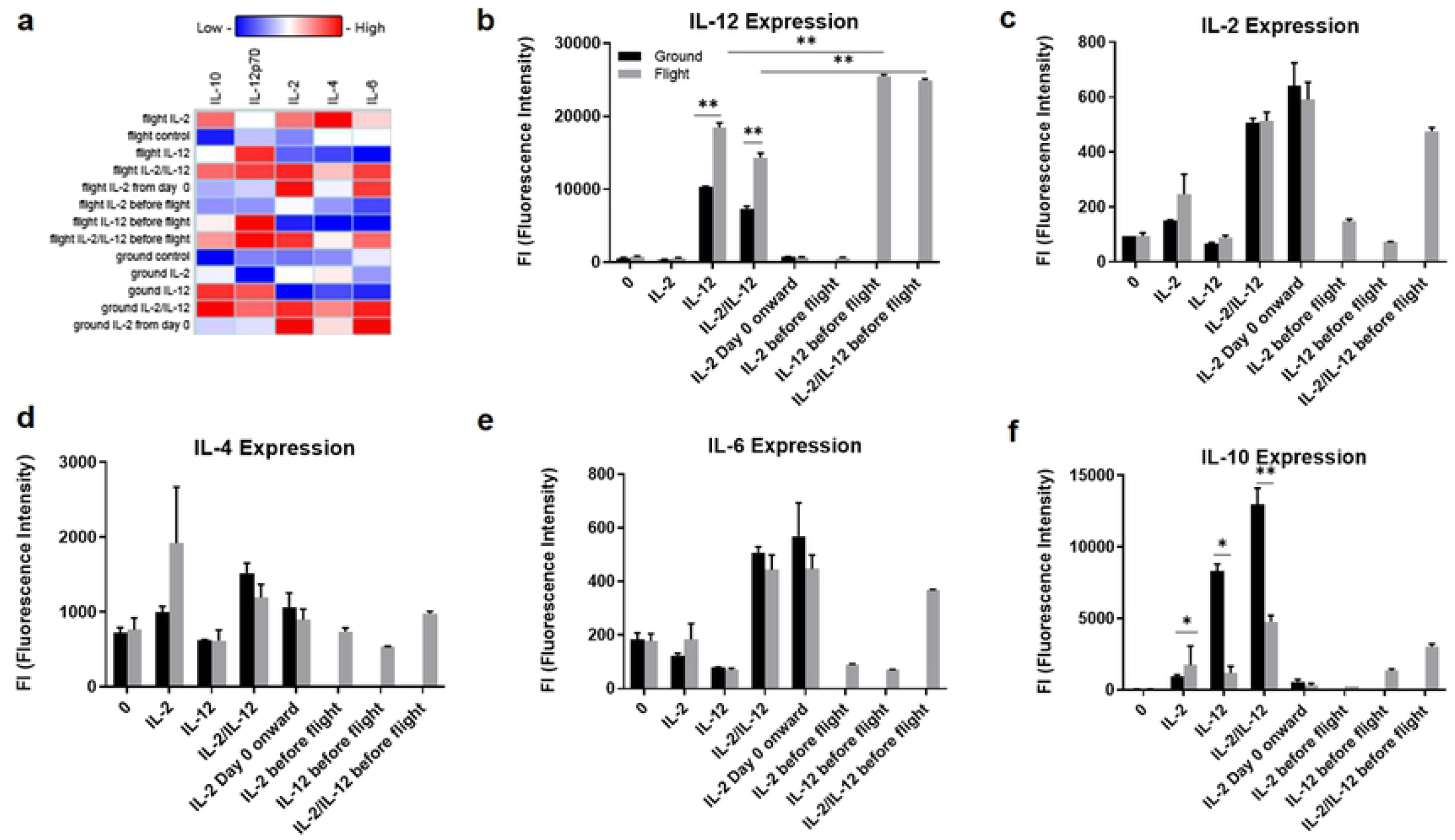
Exposure to microgravity alters the expression of multiple cytokines detected from T cell culture supernatants. Supernatants of flown cells and grounds controls were collected post flight and assessed for IL-12, IL-2, IL-4, IL-6, and IL-10 expression (fluorescence intensity) by multiplex immunoassay. (a) Indicates the heat map of all conditions for all cytokines. (b) Shows IL-12 expression, (c) indicates IL-2 expression, (d) IL-4 expression, (e) IL-6 expression, and (f) IL-10 expression. Different y axis scales used to demonstrate the profound differences across different conditions. Data represented as means ±SEM. ** p<0.01, * p<0.05. Significance was determined by using t-tests. Black color indicates ground conditions, grey shows flight conditions.

Specifically, our data show that the average value for each group (IL-2, IL-12, IL-4, IL-6, and IL-10) is 81.39%, 57.57%, 27.31%, 19.13%, and 54.86%, respectively. Another interesting observation is that the % CV (control) is the largest for IL-2 day 0 for each block of IL-2, IL-4, IL-6 and IL-10. Similarly, the % CV (flight) shows one of the largest values for IL-2 across each block in Table 3. We observed a significantly enhanced expression of IL-12 in flown T cells that were cultured in IL-12 alone (p < 0.01) or in combination with IL-2 (p < 0.01) as compared to ground controls (Fig 6B). This increase in IL-12 is consistent with the functional data which indicated the elevated expression of IFNɣ in T cells cultured in IL-12. Adding IL-12 before the flight increased the expression of IL-12 in T cells cultured in IL-12 alone or in combination with IL-2. However, cells exposed to suborbital flight and cultured in IL-2 from day 0 had a lower expression of IL-12 (Fig 6B). Analysis of IL-2 expression revealed that this cytokine was slightly elevated in all flown T cell conditions, except for cells cultured in IL-2 from day 0 when compared to ground controls (Fig 6C). However, the difference in expression was not significant among different ground and flight conditions. The same pattern was observed in IL-4 expression, except for T cells cultured with IL-2/IL-12, which when exposed to suborbital flight had a lower expression of IL-4 as compared to ground controls (Fig 6D). The expression of cytokine IL-6 was reduced in all flight conditions except for the cells cultured in IL-2 (Fig 6E). Interleukin-6 is a pleiotropic cytokine mediating and facilitating the inflammatory responses. Literature suggests that IL-6 might promote T cell survival and proliferation [33, 34], although the precise mechanisms accounting for these effects are not well-delineated. Our data indicate that culturing T cells with IL-2/IL-12 and IL-2 from day 0 leads to the highest expression of IL-6 across all conditions, but with lower expression in flight conditions, relative to ground controls. Given IL-6 may drive T cell survival, our data is in alignment with our other findings suggesting that microgravity may have a negative effect on T cell survival and proliferation. Finally, we saw a significant higher expression (p < 0.05) of IL-10 in flown T cells cultured in IL-2, but significantly reduced expression in flown cells supplemented with either IL-12 or both cytokines (Fig 6F). IL-10 is classified as an anti-inflammation marker involved in antibody production and regulation of inflammation. However, its role in modulating the immunological response might be quite complicated [35]. Our data show that microgravity significantly reduced the expression of IL-10, however culturing T cells in IL-12 may restore the expression of IL-10 to some degree, but not fully. Together, our data demonstrate that multiplex immunoassay is a sensitive assay capable of detecting different analytes in T cell cultures, exposed to microgravity. Furthermore, adding cytokines to T cell cultures may restore some of the antagonizing effects of microgravity, such as an increase expression of IL-12 that has the ability to stimulate both innate and adaptive immune response and may play a role in the development of therapeutic modalities to repair immune dysfunctions.

## Conclusions and discussion

The primary goal of this study was to investigate whether brief exposure to microgravity (3.2 min) during suborbital space flight could affect the behavior and function of the activated murine T cells. Furthermore, our secondary objective was to determine whether culturing cells with cytokines IL-2 and IL-12 alone or in synergistic manner could reverse the suggestive antagonizing effect of microgravity.

To show the microgravity induced T cell behavior alterations, we assayed several parameters, including T cells activation markers, cell viability and apoptosis, functional responsiveness to stimulants and cytokine secretion abilities. Overall, our data demonstrated that brief exposure to microgravity affected T cell behavior by altering CD8+ and CD4+ expression, reducing functional capabilities and inducing apoptosis in some conditions, but not in all. Given such a brief exposure to microgravity (3.2 minutes) and seeing a reduction of many parameters in flight conditions compared to ground controls, we can conclude that T cells might be sensitive to microgravity even at this short exposure. However, a much longer time exposure would be needed to observe a more robust effect. Culturing T cells with different cytokines exhibited differences in their responses to microgravity. As such, our data revealed that exposure to suborbital flight reduced the expression of CD4/CD8 ratio, with the highest difference observed in cells cultured with IL-2 from day 0. This could be explained by the IL-2 receptor signaling and its higher responsiveness to CD8 than to CD4 [36]. Also, our data show that brief exposure to microgravity alters the expression of activation markers CD69, CD71, and CD25 with the highest downregulation effect observed in CD71 expression. Supplementing T cells with IL-2 and IL-12 increased the expression of CD71. Also, our findings suggest that adding IL-12 to T cell cultures restores the expression of CD69. Consistent with previous reports, our study also demonstrate that microgravity might shift cells into more apoptotic phase, but cytokines could rescue them from dying, especially IL-2. The benefit of adding cytokines to T cell cultures were also demonstrated in increased functional capabilities. Finally, our study also provides additional insights on cytokine expression profiles derived from flown T cell supernatants and their matched ground controls. Also, it is important to note that due to limited weight allowance of the suborbital payload, some of our cells were not cultured in duplicate and therefore the difference observed between ground and flight conditions might have failed to reach significance. Taken together, our results demonstrate that even a brief exposure of about 3.2 minutes to microgravity leads to some alterations in cellular responses of T cells. Thus, our data suggest that adding IL-2 and IL-12 is a simple strategy to mitigate the downregulated effect of microgravity on some T cell cellular responses, but not in all. Our current work will be further tested using other space research platforms, such as the PLD Space’s Miura 1 rocket in 2022-2023.

In general, the current knowledge of microgravity induced alterations in the immune system derives from multiple sources, such as leukocytes obtained from astronauts, cells obtained from flown animals (mostly mice and rats), and from in-vitro cell cultures. Therefore, comparison across these different model systems should be made with caution. Although it is quite difficult to compare our study’s results with other studies due to different microgravity subjects and models used and different assessment time points, our data supports findings from previous studies on microgravity induced alterations on the immune system. However, given that we manipulated T cells by culturing them in different cytokines, we supplemented current knowledge by providing additional insights on how T cells, derived from different culturing conditions, respond to microgravity. We cannot rule out the possibility that our results might be influenced by the temperature changes among flight and control conditions, limited number of samples, or inability to fix our sample after microgravity phase in flight (samples returned about 5 hours after launch). However, we attempted to mimic the physiological conditions of flown T cells to ground controls to the best of our abilities. More future studies are needed to delineate the mechanisms mediating microgravity effect on immunity. Thus, based on the most comprehensive experiment conducted in space, the NASA Twin Study, the time of the cytokine expression assessment could also play a role [37]. Therefore, future studies should aim to assess the behavior of T cell prior to, during and post flight time points. The exposure of T cells (non-activated human Jurkat T lymphocytic cells) to microgravity during suborbital flight was assessed before as part of the German Aerospace Center (DLR) gravitational biology experiment on a suborbital rocket (TEXUS-51) in 2017. T cells were subjected to vibrations and hypergravity levels and demonstrated the high levels of stability of gene expression [21]. However, more studies are needed to investigate the effect of microgravity on T cells behavior during the suborbital flight so that acquired data could be used to design comprehensive orbital space platform experiments. Our team hopes to replicate our findings on the next suborbital flight onboard the PLD Space’s Miura 1 rocket where we will extend our analysis to gene expression and signal transduction assays before moving on to orbital research platform. One of the major concerns of long space missions is a risk to develop cancer [38]. Due to limited data available, it is hard to estimate such risk for future space travelers. However, knowing that microgravity does induce immune downregulation which, coupled with radiation, alterations in circadian rhymes and other side effects, could place humans at a high risk to develop malignant diseases. Therefore, more studies are needed to explore the role of different cytokines known to modulate the immune response to microgravity. There are already technologies developed in space, such as microencapsulation for the new drug delivery system and light technology for the pain relief applications which assist in cancer therapies on Earth [8]. Thus, given that space environment is challenging and characterized by symptoms similar to those experienced by cancer patients (such as fatigue, disruption of circadian rhymes), strategies to counteract these effects should be designed. The unique environment of microgravity coupled with advances in laboratory techniques serves a dual purpose: these help to gain insights into T cells behavior which in turn could help to design interventions to enhance the quality of life for those travelling to space and battling cancer on Earth.

## Acknowledgements

We would like to thank the Applied Aviation Sciences Department of the College of Aviation at the Embry Riddle Aeronautical University (ERAU) for their partial support to Dr. Pedro Llanos during his stay during the summer of 2016 to initiate this research at the Medical University of South Carolina (MUSC). Thanks to the Payload Applied Technology and Operations (PATO) lab in the College of Aviation, and the College of Engineering, to the Mechanical Engineering Department for providing laboratory its resources for ground testing. In addition, we thank Dr. Mark Rubinstein for providing antibodies and other reagents required for preparing T cells and for the expert advice in culturing murine T cells during the various feasibility studies conducted in 2016-2017 prior to suborbital launch. Also, we would like to thank Dr. Mike Wargovich’s lab for providing resources and tools to prepare and assess T cells during pre- and post-flight operations. Finally, authors would like to thank Leonid Bunegin for his help with the life support system to house T cells during transportation to the launch site.

## Contributions

K.A. and P.J.L. designed and performed feasibility and experimental studies and wrote the manuscript. P.J.L. and S.G. and V.D. conceptualized, tested, and supervised the final design of the hardware where the cells were housed to meet the NanoRacks milestones. J.M. helped performing and analyzing the cytokine expression assay with the Bio-Plex kit provided by the P.J.L.’s FIRST grant. M.W. assisted in providing resources to conduct pre- and post-launch experiments and data analyses. All authors reviewed and edited the manuscript.

